# Acute fluoxetine differently affects aggressive display in zebrafish phenotypes

**DOI:** 10.1101/217810

**Authors:** Hellen Pereira Barbosa, Monica Gomes Lima, Caio Maximino

## Abstract

Zebrafish have been introduced as a model organism in behavioral neuroscience and biological psychiatry, increasing the breadth of findings using fish to study the neurobiology of aggression. Phenotypic differences between leopard and longfin zebrafish were exploited in order to elucidate the role of phasic serotonin in aggressive displays on this species. The present study revealed differences in aggressive display between leopard and longfin zebrafish, and a discrepant effect of acute fluoxetine in both populations. In mirror-induced aggression, leopard animals showed higher display latencies than longfin, as well as lower display duration and frequency (Experiment 1). Moreover, 2.5 mg/kg fluoxetine decreased the duration and frequency of display in longfin, but not leopard; and 5 mg/kg fluoxetine increased display frequency in leopard, but not longfin (Experiment 2). It is suggested that zebrafish from the longfin phenotype show more aggressive motivation and readiness in the mirror-induced aggression test that leopard, and that acute fluoxetine increases aggression in leopard and decreased it in longfin zebrafish.

## 1. Introduction

The biological comprehension of factors underlying aggression is still limited (Miczek et al., 2007), even though a range of mental disorders present aggression as a symptom (Krakowski, Volavka, & Brizer, 1986). Despite the paucity of neurobiological data on aggression, a role for monoamines has been proposed (Miczek et al., 2007; Takahashi, Quadros, Almeida, & Miczek, 2011). In most animal models, acutely increasing the serotonergic transmission inhibits aggressive behavior (Takahashi et al., 2011); a metanalysis of preclinical studies demonstrated that, across species, pharmacologically increasing 5-HT levels inhibits aggression (Carrillo, Ricci, Coppersmith, & Melloni Jr., 2009).

While this observation appears to hold for most studies, some controversies and gaps appear in the literature, especially in basal vertebrates such as fish. For example, in the metanalysis by Carrillo et al. (2009), 5-HT decreased aggression in wrasses and trouts, but not in the Siamese fighting fish. Zebrafish have been introduced as a model organism in behavioral neuroscience and biological psychiatry (Norton & Bally-Cuif, 2010; Stewart et al., 2015), increasing the relevance of findings using fish to study the neurobiology of aggression.

A role for 5-HT in zebrafish aggressive behavior has been suggested by neurochemical studies. After eliciting an aggressive display towards a mirror (mirror-induced aggression, MIA), 5-HT levels were increased in the telencephalon, while 5-HIAA was increased in the optic tectum of zebrafish (Teles, Dahlbom, Winberg, & Oliveira, 2013). Male and female zebrafish respond to agonistic encounters in a similar fashion; nonetheless, males present higher 5-HT turnover in the forebrain in relation to females, suggesting that aggressive bouts could be more stressful to males than females (Dahlbom, Backström, Lundstedt-Enkel, & Winberg, 2012). Filby et al. (2010) demonstrated that dominant males show an overexpression of genes associated with the serotonergic system in the hypothalamus, including *tph1b* and *htr1aa*, while females showed overexpression of *tph2*, *htr1aa*, *slc6a4a*, and *mao* in the hypothalamus and *tph1a* and *tph2* in the telencephalon.

While these results suggest that aggressive behavior can be linked to differences in the serotonergic system – especially in the context of dominance hierarchies –, a causal relationship is more tenuous. Filby et al. (2010) treated dominant male zebrafish with fluoxetine (3 or 4.5 µg/L), without effects on aggressive behavior in a dyadic encounter; however, a similar concentration (3 µg/L) decreased aggressive displays in the MIA (W. H. J. Norton et al., 2011). Using a much higher concentration (5 mg/L), Theodoridi et al. (2017) were able to inhibit attacks and chasing behavior in dominant animals in dyads. The lack of consistency could be due to dosing, behavioral paradigms (e.g., MIA vs. dyadic encounters), or other variables.

Recent studies also showed that 5-HT levels are lower in zebrafish with the leopard phenotype than in animals with the longfin phenotype, an alteration that is accompanied by increased monoamine oxidase activity (Maximino, Puty, Oliveira, & Herculano, 2013). These neurochemical differences were accompanied by increased anxiety-like behavior that is rescued by fluoxetine treatment (Maximino, Puty, Oliveira, et al., 2013). Interestingly, in longfin animals fluoxetine *increases* anxiety, and 5-HT levels are negatively correlated with anxiety-like behavior; it is possible that embryological differences in the serotonergic system produce opposite adult phenotypes.

These phenotypic differences are exploited in the present work to clarify the role of phasic serotonin on aggressive displays in zebrafish. We hypothesized that the hyposerotonergic phenotype of leopard zebrafish would produce increased aggressive behavior, and that fluoxetine would rescue this phenotype. The experimental evidence produced in the present work contradicted this hypothesis, since longfin were shown to display more aggressive motivation and readiness in the mirror-induced aggression test than leopard zebrafish, and since acute fluoxetine increased aggression in leopard animals and decreased it in longfin zebrafish. This manuscript is a complete report of all the studies performed to test the effect of skin phenotype and fluoxetine on aggressive behavior. We report how we determined our sample size, all data exclusions (if any), all manipulations, and all measures in the study.

## 2. Methods

### 2.1. Animals, housing, and baseline characteristics

Outbred populations were used due to their increased genetic variability, decreasing the effects of random genetic drift which could lead to the development of uniquely heritable traits (Parra, Adrian Jr, & Gerlai, 2009; Speedie & Gerlai, 2008). Thus, the animals used in the experiments are expected to better represent the natural populations in the wild. Adult zebrafish from the wildtype strain (longfin and leopard phenotypes) were used in this experiment. Animals were bought from a commercial vendor, and arrived in the laboratory with an approximate age of 3 months (standard length = 13.2 ± 1.4 mm), and were quarantined for two weeks; the experiment began when animals had an approximate age of 4 months (standard length = 23.0 ± 3.2 mm). Animals were kept in mixed-sex tanks during acclimation, with an approximate ratio of 50 male:50 female. Both phenotypes were kept in the same tank before experiments. The breeder was licensed for aquaculture under Ibama’s (Instituto Brasileiro do Meio Ambiente e dos Recursos Naturais Renováveis) Resolution 95/1993. Animals were group-housed in 40 L tanks, with a maximum density of 25 fish per tank, for at least 2 weeks before experiments begun. Tanks were filled with non-chlorinated water at room temperature (28 °C) and a pH of 7.0-8.0. Lighting was provided by fluorescent lamps in a cycle of 14-10 hours (LD), according to standards of care for zebrafish (Lawrence, 2007). Water quality parameters were as follows: pH 7.0-8.0; hardness 100-150 mg/L CaCO3; dissolved oxygen 7.5-8.0 mg/L; ammonia and nitrite < 0.001 ppm. All manipulations minimized their potential suffering of animals, and followed Brazilian legislation (Conselho Nacional de Controle de Experimentação Animal - CONCEA, 2017). Animals were used for only one experiment and in a single behavioral test, to reduce interference from apparatus exposure.

### 2.2. Mirror-induced aggression

The protocol for mirror-induced aggression was adapted from Norton et al. (2011). Animals were individually transferred to a tank (15 × 10 × 30 cm) containing 1 L of holding tank water. The tank was lit from above with white light (445 ± 56 lumens). Tanks were not aerated during testing, so as not to disturb the animals. However, system water, which shows adequate D.O. levels (7.5-8.0 mg/L), was used thoughout the experiments. Animals were allowed to acclimate to the tank for 5 min; after that, a mirror was positioned on the outside of the tank (on the narrower side), in an angle of 22.5º. All steps of the experiment were executed under constant Gaussian white noise (55 ± 2.5 dB above the tank). Behavior was recorded with a digital video camera (Samsung ES68) positioned in the wider side of the tank, and analyzed by observers blind to treatment using the event-recording software X-Plo-Rat (https://github.com/lanec-unifesspa/x-plo-rat). The following endpoints were analyzed: time in the square nearest to the mirror (s); frequency (N) and duration (s) of aggressive display; total number of squares crossed (N). Aggressive display was defined as a swimming posture with erect dorsal, caudal, pectoral, and anal fins (Gerlai, Lahav, Guo, & Rosenthal, 2000).

### 2.3. Data availability

Datasets and scripts for all analyses are available from https://github.com/lanec-unifesspa/5HT-aggression (doi: 10.5281/zenodo.1006701).

### 2.4. Experiment 1

#### Sample size calculation and groups

Sample sizes were calculated on results regarding the effects of fluoxetine on total time in broadside display in *Betta splendens*, reported by Lynn et al. (2007); as a result, the closest endpoint (time on display) was chosen as the primary endpoint, and calculations for sample sizes are valid only for that endpoint. Calculations were based on Rosner’s (2016) method for comparing two means, and assumed α = 0.05 and power 80% on a two-tailed analysis. Based on these calculations, 15 animals were used in each group in Experiment 1. Animals were derived from the stock population described in section 2.1, and displayed either the longfin (Group LOF) or leopard (Group LEO) phenotypes. Animals were randomly drawn from the tank immediately before testing, and the order with which phenotypes were tested was randomized *via* generation of random numbers using the randomization tool in http://www.randomization.com/. Blinding was not possible, due to the obvious differences in skin phenotype.

#### Experimental design and statistical analysis

Animals were allocated to each group according to phenotype. Immediately after being drawn from the tank, animals were individually transported to the experiment room, and left undisturbed for 30 min. After this interval, animals were exposed to the MIA test, described above. Differences between groups were analyzed using Approximative Two-Sample Fisher-Pitman Permutation Tests 10,000 Monte-Carlo re-samplings, using the R package ‘coin’ (Hothorn, Hornik, van de Wiel, & Zeileis, 2006). The data analyst was blinded to phenotype by using coding to reflect treatments in the resulting datasets; after analysis, data was unblinded. Data are presented using individual dot plots combined with boxplots. Effect sizes are noted in the text as Cohen’s d.

### 2.5. Experiment 2

#### Sample size calculation and groups

In the absence of similar experiments in the literature, sample sizes were calculated based on the assumption of fixed effect sizes for both the phenotype and the dose factors, with a projected effect size of 0.4, and 80% power; calculations were made using the R package ‘pwr2’ (Lu, Liu, & Koestler, 2017). Based on these calculations, 10 animals were used in each group in Experiment 1. Animals were derived from the stock population described in section 2.1, and displayed either the longfin (Group LOF) or leopard (Group LEO) phenotypes. Animals were randomly drawn from the tank immediately before testing, and the order with which phenotypes were tested was randomized *via* generation of random numbers using the randomization tool in http://www.randomization.com/. Blinding for phenotype was not possible, due to the obvious differences in skin phenotype. Animals from each phenotype were randomly allocated to treatment (vehicle or either fluoxetine dose) *via* generation of random numbers using the randomization tool in http://www.randomization.com/.

#### Drug treatments

Fluoxetine (FLX) was bought from EMS, dissolved in Cortland’s salt solution (Wolf, 1963), and injected intraperitoneally in cold-anesthetised animals (Kinkel, Eames, Philipson, & Prince, 2010). FLX doses (2.5 and 5.0 mg/kg) were based on the demonstration of effect on the light/dark test on longfin (Maximino, Puty, Benzecry, et al., 2013) and leopard (Maximino, Puty, Oliveira, et al., 2013) zebrafish. Experimenters were blinded to treatment by coding drug vials.

#### Experimental design and statistical analysis

Animals were allocated to each group according to phenotype. Immediately after being drawn from the tank, animals were individually transported to the experiment room, injected with vehicle or drug, and left undisturbed for 30 min. After this interval, animals were exposed to the MIA test, described above. Differences between groups were analyzed using two-way analyses of variance with robust estimators on Huber’s M-estimators, using the R package ‘rcompanion’ (Mangiafico, 2017). P-values were adjusted for the false discovery rate. The data analyst was blinded to phenotype by using coding to reflect treatments in the resulting datasets; after analysis, data was unblinded. Data are presented using individual dot plots combined with boxplots. Effect sizes are reported as partial ε² values, and were calculated using the R package ‘lsr’ (Navarro, 2015).

## 3. Results

### 3.1. Experiment 1

LEO zebrafish showed longer latencies to display than LOF animals (Z = 3.3925, p = 0.0005, *d* = 1.5779; Figure 1A), as well as shorter display durations (Z = −2.5659, p = < 2.2e-16, *d* = −1.0605; Figure 1B) and frequency (Z = −2.7073, p = 0.003, *d* = −1.1372; Figure 1C), and time spent near the mirror (Z = −3.2284, p = < 2.2e-16, *d* = −1.4593; Figure 1D). No differences were found in total locomotion (Z = −0.69887, p = 0.4965, *d* = −0.2573; Figure 1E).

**Figure 1.**
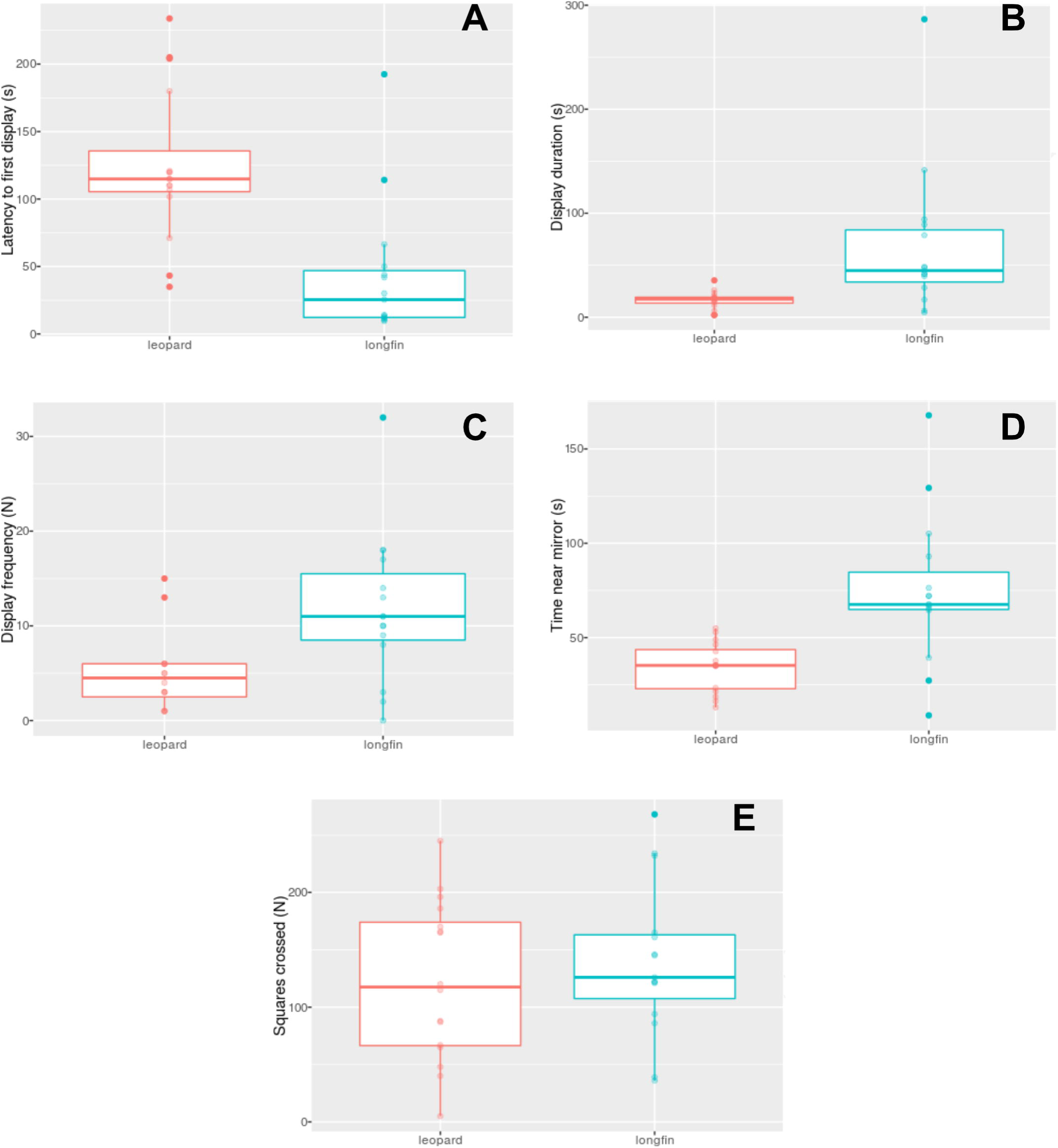
Phenotype differences in (A) latency to display, in s; (B) display duration, in s; (C), display frequency; (D), time spent near the mirror, in s; and (E), total locomotion. Boxplots represent median and interquartile range, with Tukey whiskers.

### 3.2. Experiment 2

No main effects of phenotype (p = 0.5736; partial ε² = 0.0326) or FLX dose (p = 0.5044; partial ε² = 0.0482) were found for latency, but a significant interaction was found (p = 0.0412; partial ε² = 0.1622); nonetheless, post-hoc tests did not detect any differences between groups (Figure 2A). Main effects of phenotype (p = 0.0132; partial ε² = 0.2429) and FLX dose (p = 0.0062; partial ε² = 0.2297), as well as an interaction effect (p = 0.009; partial ε² = 0.1986), were found for display duration (Figure 2B). Post-hoc tests suggested that FLX (2.5 mg/kg) decreased display duration on LOF, but not LEO (p = 0.032 vs. 0 mg/kg). Similarly, main effects of phenotype (p = 0.0458; partial ε² = 0.2744) and FLX dose (p = 0.004; partial ε² = 0.2476), as well as an interaction effect (p < 0.0001; partial ε² = 0.3598), were found for display frequency (Figure 2C); post-hoc tests suggested that FLX (5 mg/kg) increased display frequency in LEO, but not LOF animals (p = 0.004 vs. 0 mg/kg). No main effects of phenotype (p = 0.2692; partial ε² = 0.3704) were found for time near mirror, but a main effect of FLX dose (p = 0.0004; partial ε² = 0.0164) and an interaction effect (p < 0.0001; partial ε² = 0.5760); post-hoc tests suggested a inverted-U-shaped curve for FLX-treated LOF (0 vs. 2.5 mg/kg: p = 0.0012; 0 vs. 5.0 mg/kg: p = 0.0167), while a monotonic increase for FLX-treated LEO (0 vs. 2.5 mg/kg: p = 0.0496; 0 vs. 5.0 mg/kg: p < 0.0001) (Figure 2D). No main or interaction effects were found for total locomotion (Figure 2E).

**Figure 2.**
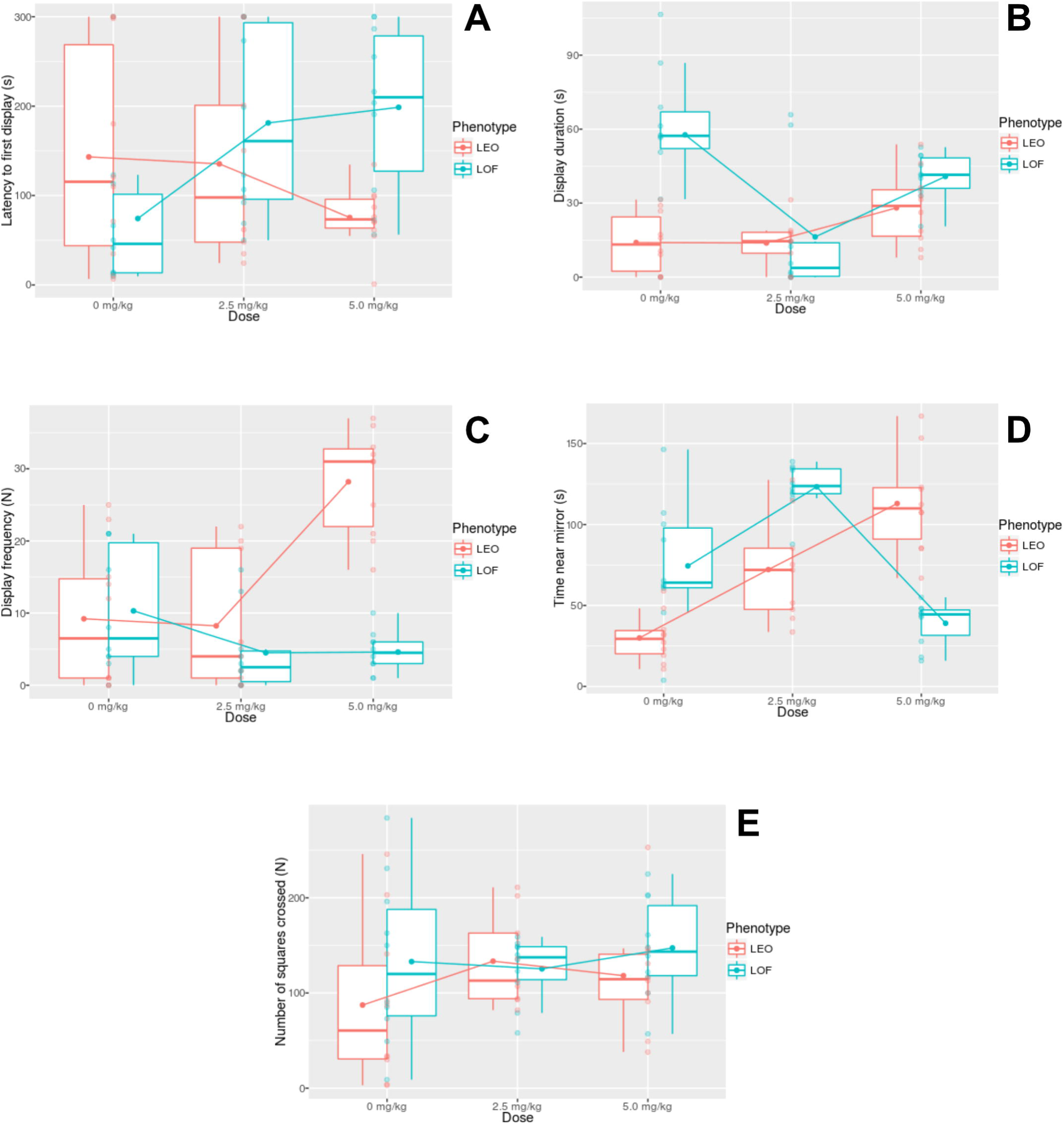
Effects of fluoxetine on the aggressive display of longfin (LOF, dark gray) and leopard (LEO, light gray) zebrafish. (A) latency to display, in s; (B) display duration, in s; (C), display frequency; (D), time spent near the mirror, in s; and (E), total locomotion. Boxplots represent median and interquartile range, with Tukey whiskers. Dots joined by lines represent means.

## 4. Discussion

The present work demonstrated that zebrafish with the leopard skin phenotype show less aggressive readiness and less aggression in relation to longfin animals. Moreover, a different pattern of fluoxetine effects was observed, with fluoxetine decreasing aggressive display (but not readiness) in longfin animals and increasing it in leopard animals. Evidence for dose-dependence was also observed.

Serotonin (5-HT) has long been implicated in the neurobiological mechanisms of aggressive behavior (Miczek et al., 2007; Summers & Winberg, 2006; Takahashi et al., 2011). In a metanalysis of preclinical studies, Carrillo et al. (2009) demonstrated that, across species, pharmacologically increasing 5-HT levels inhibit aggression. Interestingly, in their metanalysis a species-specific effect was found in fish, with 5-HT decreasing aggression in wrasses and trouts, but not in the Siamese fighting fish (Carrillo et al., 2009). The present study examined only aggressive displays, which was elicited in the mirror-induced aggression test; as a result, dominance hierarchies were not induced. Similar results were observed by Norton et al. (2011), which observed reduced aggressive displays in Tübingen zebrafish treated with fluoxetine. When zebrafish are allowed to form dominance hierarchies, fluoxetine either produces no effect (Filby et al., 2010, using WIK zebrafish) or reduces aggression in dominant males, but not in subordinates (Theodoridi et al., 2017, undescribed phenotype). Given that these studies used different phenotypes than those reported here, conclusions are limited.

Behavioral differences between leopard and longfin phenotypes were observed in zebrafish before (Canzian, Fontana, Quadros, & Rosemberg, 2017; Egan et al., 2008; Maximino, Puty, Oliveira, et al., 2013; Quadros et al., 2016; https://doi.org/10.1101/055657). Of special relevance is the observation that leopard zebrafish present increased brain monoamine oxidase (MAO) activity that is associated with lower serotonin levels and higher turnover of 5-HT in the brain (Maximino, Puty, Oliveira, et al., 2013; Quadros et al., 2018). This hyposerotonergic profile was also associated with increased anxiety-like behavior that was rescued by fluoxetine treatment (Maximino, Puty, Oliveira, et al., 2013). Moreover, Quadros et al. (2018) also found that leopard to be less aggressive than shortfin zebrafish, suggesting a consistent hypoaggressive phenotype across laboratories, conditions, and background genetics.

These results suggest a serotonin-linked behavioral syndrome in zebrafish that varies across populations. Indeed, the hyperanxious profile observed in leopard (Maximino et al., 2013) is also rescued by fluoxetine treatment at the same dose range as that reported here. A variety of studies in Siamese fighting fish (*Betta splendens*) suggest that fluoxetine reduces aggressive behavior (Dzieweczynski & Hebert, 2012; Eisenreich & Szalda-Petree, 2015; Kania, Gralak, & Wielgosz, 2012) and boldness (Dzieweczynski, Campbell, & Kane, 2016; Dzieweczynski, Kane, Campbell, & Lavin, 2016), which could be interpreted as either increased impulsivity or decreased anxiety. The presence of a aggression-boldness syndrome has long been proposed in as a dimension in fish behavior (Conrad, Weinersmith, Brodin, & Saltz, 2011), and the literature appears to point to serotonin as an important link in that. Nonetheless, these results must be interpreted with caution, given that the *B. splendens* experiments were made with chronic fluoxetine treatment (which, along with increased serotonin levels, is thought to induce other long-term neuroadaptations; Castrén & Antila, 2017). Moreover, other experiments with zebrafish (Norton et al., 2010) failed to find an effect of fluoxetine in aggression-boldness – although, again, the use of different strains and phenotypes make it difficult to generalize.

The results from the Carrillo et al. (2009) metanalysis suggested a inhibitory role for phasic serotonin (i.e., 5-HT released by either the aggressive act itself, or by pharmacological manipulations such as fluoxetine); a role for tonic 5-HT, in *Betta splendens*, was discarded, because neither the 5-HT synthesis inhibitor *para*-chlorophenylalanine nor the 5-HT precursor L-tryptophan changed display behavior (Clotfelter, O’Hare, McNitt, Carpenter, & Summers, 2007). If that was also true for zebrafish, differences in aggressive display between leopard and longfin would not be expected, given that these phenotypes differ in serotonergic tone (Maximino, Puty, Oliveira, et al., 2013). While these differences were observed in the present work, they occur in the opposite direction from what would be predicted from 5-HTergic tone alone (i.e., we should expect leopard zebrafish to be more aggressive if aggression was linearly and negatively related to tone). It is more likely that the normalization of 5-HT levels in leopard after fluoxetine treatment is responsible for increased aggression, while the “extra” 5-HT levels after fluoxetine treatment in longfin reduce its basal aggression levels; as a result, the relationship between aggression and 5-HT levels are to be interpreted as following and inverted-U-shaped distribution (Figure 3). This is also reinforced by the generally hormetic dose-response curves observed in longfin animals treated with fluoxetine. Alternatively, it is possible that a developmental effect is responsible for these discrepancies.

**Figure 3.**
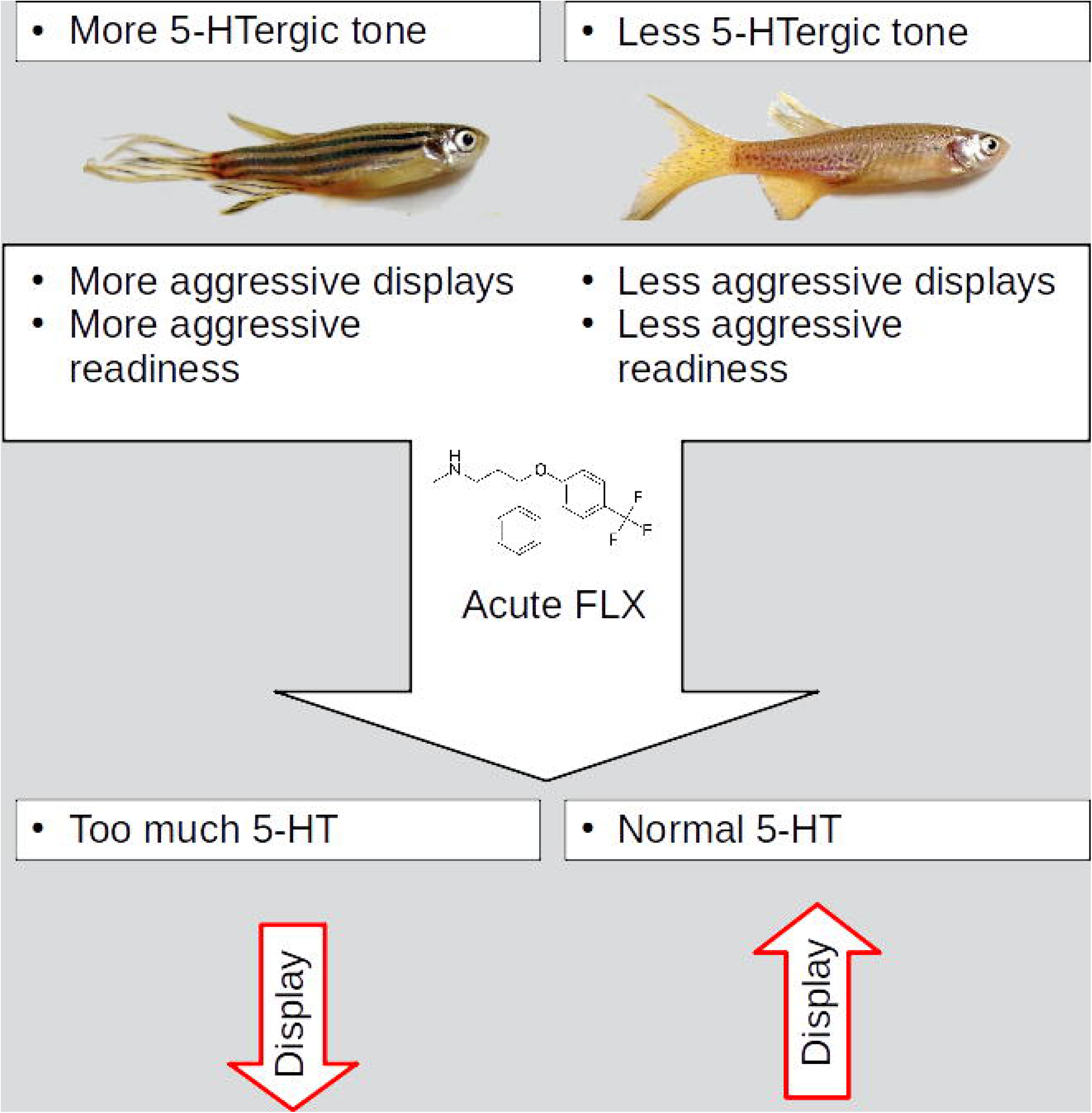
Hypothesized role of the serotonergic tone on the organization of aggressive display in longfin and leopard zebrafish.

While it might be tempting to attribute these differences to genetic differences across populations, the animals used were not derived from inbred strains. The altered pigmentation observed in our leopard fish has previously been reported, in the Tupfel long-fin (TL) strain, to be due to a mutation in *connexin41.8* (Watanabe et al., 2006); nonetheless, it is unknown whether this mutation is present in our animals, or whether this genetic marker is one of many loci that differ between leopard and longfin animals (Gerlai, 2018). These differences make it difficult to make specific genetic inferences regarding the differences observed in the present work. Nonetheless, behavioral differences between leopard and longfin zebrafish were consistently observed across laboratories (and therefore across fish vendors) (Canzian, Fontana, Quadros, & Rosemberg, 2017; Egan et al., 2008; Maximino et al., 2013; Quadros et al., 2016, 2018), and the effect of phenotype on zMAO activity was observed independently at least twice (Maximino et al., 2013; Quadros et al., 2018), suggesting an important link between the serotonergic system and aggressive behaviors across zebrafish strains.

These results are also reminiscent of what is observed in different populations of *Astyanax mexicanus*. In that species, different populations occupy different niches, and surface-dwelling populations are much more aggressive than cave-dwelling populations (Rétaux & Elipot, 2013). These differences are related to the density of serotonergic neurons in the hypothalamus, with cavefish showing a higher number of 5-HT neurons in that region (Elipot, Hinaux, Callebert, & Rétaux, 2013). Moreover, a mutation in the *mao* gene was found in cavefish that led to an hyperserotonergic phenotype (Elipot et al., 2014). Treating surface fish with fluoxetine decreases aggression, while in cavefish the drug slightly increases it (Elipot et al., 2013). In the present paper, however, treatment with fluoxetine increased aggression in leopard animals (which show an hyposerotonergic profile in relation to longfin animals; Maximino, Puty, Oliveira, et al., 2013) and decreased it in longfin zebrafish. These differences might be due to the origin of serotonin, since, in *Astyanax mexicanus* populations, raphe 5-HT levels are unchanged, while hypothalamic 5-HT is increased (Elipot et al., 2013); further experiments are needed to untangle this hypothesis.

Interestingly, a different dose-response profile was observed between phenotypes in the present study, with the low dose (2.5 mg/kg) generally decreasing aggression in longfin and the high dose (5.0 mg/kg) generally increasing it in leopard. While difficult to explain presently, these results suggest either that an “optimal” serotonergic tone is needed to maintain aggression levels, or that serotonin transporters are desensitized or downregulated in the leopard population. While the first hypothesis is more likely, given the observation of an hyposerotonergic profile in leopard zebrafish (Maximino et al., 2013; Quadros et al., 2018), the current state of the literature and the current data are not enough to assess this.

In conclusion, the present experiments revealed differences in aggressive behavior between leopard and longfin zebrafish, and a discrepant effect of fluoxetine on both populations. These results are relevant to understand the role of tonic and phasic serotonin neurotransmission on aggressive behavior in preclinical models, and might contribute to a better appreciation of the complex roles of this monoamine in controlling vertebrate aggression.

## Acknowledgments

Data packages and statistical analysis scripts for this article can be found at https://dx.doi.org/10.5281/zenodo.1006701.

